# Unbiased proteomic analysis detects painful systemic inflammatory profile in the serum of nerve injured mice

**DOI:** 10.1101/2022.02.28.482355

**Authors:** Wen Bo Sam Zhou, Xiang Qun Shi, Younan Liu, Simon D. Tran, Francis Beaudry, Ji Zhang

**Author notes:** <u>Corresponding authors:</u> Dr. Ji Zhang, Alan Edwards Centre for Research on Pain, McGill University, 740 Docteur Penfield Ave, Suite 3200C, Montreal, QC, Canada H3A0G1.

## Abstract

Neuropathic pain is a complex, debilitating disease that results from injury to the somatosensory nervous system. The presence of systemic chronic inflammation has been observed in chronic pain patients, but whether it plays a causative role remains unclear. This study aims to determine the perturbation of systemic homeostasis by an injury to peripheral nerve and its involvement in neuropathic pain. We assessed the proteomic profile in the serum of mice at 1-day and 1-month following partial sciatic nerve injury (PSNL) or sham surgery. We also assessed mouse mechanical and cold sensitivity in naïve mice after receiving intravenous administration of serum from PSNL or sham mice. Mass spectrometry-based proteomic analysis revealed that PSNL resulted in a long-lasting alteration of serum proteome, where the majority of differentially expressed proteins were in inflammation related pathways, involving cytokines/chemokines, autoantibodies and complement factors. While transferring sham serum to naïve mice did not change their pain sensitivity, PSNL serum significantly lowered mechanical thresholds and induced cold hypersensitivity in naïve mice. With broad anti-inflammatory properties, bone marrow cell extracts (BMCE) not only partially restored serum proteomic homeostasis, but also significantly ameliorated PSNL-induced mechanical allodynia, and serum from BMCE-treated PSNL mice no longer induced hypersensitivity in naïve mice. These findings clearly demonstrate that nerve injury has a long-lasting impact on systemic homeostasis, and nerve injury associated systemic inflammation contributes to the development of neuropathic pain.

## Introduction

Neuropathic pain is a highly debilitating condition, usually caused by a lesion or disease to the somatosensory system, that affects 7-10% of the general population [7; 63]. Although, our understanding of the underlying mechanisms leading to neuropathic pain has made revolutionary progess in the recent twenty years, current treatment options are far from satisfactory [3; 16]. A large amount of preclinical evidence strongly support the concept that neurons are not the only players in the pathology. Many neighboring non-neuronal cells, such as microglia and astrocytes in the central nervous system (CNS), satellite cells in the dorsal root ganglia (DRG), and Schwann cells in the peripheral nervous system (PNS) are actively involved [32]. The engagement of endothelial cells has also been documented [12; 35; 44]. In addition, circulating immune cells, including mast cells, neutrophils, macrophages, have been found accumulating at the injury sites. While non-neuronal cells are not expected to directly transmit electrical potentials along the pain transmission pathway, they are important modulators for neuronal activities. Upon activation, various types of molecules are released from these non-neuronal cells, facilitating cell-to-cell crosstalk. The most stunning findings are that non-neuronal cells contribute to the development and maintenance of neuropathic pain by promoting the interaction between the nervous system and the immune system [10; 33; 52].

Following the discovery on the role of neuroinflammation in neuropathic pain from animal studies, several clinical studies also revealed evidence of neuroinflammation in chronic pain patients, such as detection of inflammatory mediators in the cerebrospinal fluid (CSF) and CNS glial cell activation [2; 47]. However, due to easy access to peripheral blood samples, most studies actually reported the existence of systemic inflammation in chronic pain conditions. Some inflammatory mediators, including cytokines, chemokines have been found increased in the circulation of painful, not painless patients, associated with trauma or diabetes and in chronic widespread pain patients [2; 15; 29; 37; 38; 61]. Nevertheless, it remains unclear whether these inflammatory factors are simply correlated with chronic pain or may contribute directly to enhance pain sensitivity.

Significant advances in protein profiling technologies (protein separation, mass spectrometry, and bioinformatics) [70] offer great promise to study proteins in plasma and serum. Blood protein scanning has delivered great success in personalized health/disease assessments [18; 51; 58]. A recent in-depth proteomic study revealed undulating changes in human plasma proteome across the lifespan [39]. Another investigation of the blood samples from 16,894 participants demonstrated that plasma proteome patterns could be used as reliable and comprehensive indicators of health [71]. By taking advantage of this technology, we performed serum proteomic analysis in nerve injured mice to answer, whether and to what extent, an injury to a peripheral nerve could disturb systemic homeostasis, and whether peripheral nerve injury could lead to systemic inflammation. To further understand the functional significance of such changes in the blood to neuropathic pain, we also make use of bone marrow-derived cell extracts (BMCE), which have the potent anti-inflammatory properties [1; 59], to counteract systemic inflammation.

By combining a well-established animal model of nerve injury-induced neuropathic pain, e.g., partial sciatic nerve ligation (PSNL), high throughput serum proteomics and a functional intervention, we conclude from this study that an injury to a peripheral nerve results in a long lasting disturbance in systemic homeostasis. Serum proteins regulated by such insult are mostly involved in inflammation related pathways, indicating the existence of systemic inflammation following nerve injury. Transferring serum from nerve injured mice to naïve mice induces mechanical and cold allodynia. Reestablishing systemic balance by administration of BMCE into the blood stream of PSNL mice significantly attenuates their mechanical hypersensitivity. Serum from BMCE-treated PSNL mice no longer triggers enhanced painful response in naïve mice. Our results unravel, the potential of systemic contribution to neuropathic pain, which has been neglected in the past.

## Materials and Methods

### Animals

Experiments were performed on adult 7-9 weeks old male C57BL/6 mice, obtained from in-house breeding. Mice were housed 3 to 5 per cage in a temperature- and humidity-controlled environment with a 12/12-hour light/dark cycle, beginning at 7:00 and access to food and water ad libitum. Behavioral experiments were conducted between 9:00 and 15:00. All protocols were approved by the McGill University Animal Care and Use Committee (permit# 5533) and conducted according to the guidelines of the Canadian Council on Animal Care and the International Association for the Study of Pain. All experiments in this study using male mice, analysis for female mice will be conducted in the future.

#### 2.1 Nerve injury model

Partial sciatic nerve ligation (PSNL) was performed according to Seltzer et al. [54], adapted to mice. Briefly, mice were anesthetized with isoflurane (2.5% for induction and maintenance), and under aseptic conditions the left sciatic nerve was exposed at the thigh level. A 7-0 silk thread was used to tightly ligate approximately half of the sciatic nerve. In sham-operated animals, the nerve was exposed and manipulated, but no ligation was performed.

#### 2.2 Behavioral Analysis

Mice were habituated to the testing environment daily for at least 2 days before baseline testing. For measurements of mechanical sensitivity on the paw, withdrawal thresholds were assessed with calibrated von Frey filaments (Stoelting, Wood Dale, IL) using the up-down method [5; 9]. Mice were placed on a metal mesh floor with small Plexiglas cubicles (9 x 5 x 5-cm high) and kept for at least 1 hour for habituation before testing. A set of calibrated von Frey filaments (ranging from 0.16 g to 1.40 g of force) were applied to the plantar surface of the hindpaw until they bent. The threshold force required to elicit withdrawal of the paw (50% withdrawal threshold) was determined as the average of 2 tests separated by at least 1 hour. Mechanical sensitivity at orofacial area was also assessed by von Frey filaments as previously described [68]. Briefly, mice were placed in individual restraining cages (25 x 25 x 88 mm) and allowed to habituate for at least 1 hour before testing. A 0.04 g von Frey filament was applied perpendicularly to the orofacial area with sufficient force until it bent. One round of testing consisted of a single von Frey stimulation on one side, and a total of 20 stimulations on each side was performed per animal. A positive response was recorded if the mouse elicited a brief head withdrawal, unilateral or bilateral forepaw swipes down the snout, or an attempt to bite the von Frey filament. Results from both sides were averaged, and an increase in the total number of positive responses suggests the development of mechanical allodynia.

Cold sensitivity was evaluated with the acetone test [72]. A drop of acetone, approx. 25 μL, was applied to the plantar surface of the hind paw. The duration of acetone evoked behaviors (flinching, licking, or biting) following application of one single drop of acetone within 1 min observation was recorded. An increase in the duration of above-mentioned pain like behavior indicates cold hypersensitivity. All behavioural tests were performed blinded.

#### 2.3 Serum Collection and Transfer

A 21-gauge needle was used to puncture the facial vein of a restrained mouse, and blood was collected in Microvette 200 serum collection tubes (Sarstedt, Germany). Blood was allowed to clot for 30 minutes at room temperature before centrifugation to isolate the serum. The serum was stored at −80°C until ready for use. Serum was collected from mice at 1 day and 1 month after surgery, for both sham and PSNL groups, and two additional groups of PSNL mice received saline or BMCE treatment for 3 weeks. Serum samples were pooled from 5-6 mice/group for injections. Serum (100 μL/injection) was administered to mice intravenously through the tail vein.

#### 2.4 Proteomics analysis

The protocol for serum sample preparation was adopted from previous work [56]. Serum samples (100 μL) were denatured with 6 M urea, 20 mM dithiothreitol at 60 °C for 40 min and then alkylated at room temperature with 40 mM iodoacetamide (IAA) protected from light at room temperature for 30 min. To reduce the urea concentration, the samples were diluted 5-fold with 50 mM TRIS-HCl buffer (pH 8). Then, 5 μg of proteomic-grade trypsin was added, and the reaction was performed at 37°C for 24 h. Protein digestion was quenched by adding 10 μL of a 1% trifluoroacetic acid (TFA) solution. Samples were desalted using Hypersep Tip C18 (10-200μL) microscale solid phase extraction tips (Thermo Scientific, Rockwood, TN). Prior loading of the tryptic digest, the C18 tips were conditioned with 1 mL of methanol and then 1 mL of water containing 0.1% trifluoroacetic acid (TFA). Then the sample was loaded on the C18 tips by aspirate and dispense 10 times. The C18 tips were washed 2 times with 100 μL of 0.1% TFA and finally eluted 3 times with 100 μL of acetonitrile/water/TFA (50:50:0.1). Samples were evaporated to dryness and resuspended in 100 μL of water containing 0.1% formic acid and then transferred into injection vials for analysis.

The HPLC system was a Thermo Scientific Vanquish FLEX UHPLC system (San Jose, CA, USA). Chromatography was performed using gradient elution along with a Thermo Biobasic C18 microbore column (150 × 1 mm) with a particle size of 5 μm. The initial mobile phase conditions consisted of acetonitrile and water (both fortified with 0.1% formic acid) at a ratio of 5:95. From 0 to 3 min, the ratio was maintained at 5:95. From 3 to 123 min, a linear gradient was applied up to a ratio of 40:60, which was maintained for 3 min. The mobile phase composition ratio was then reverted to the initial conditions, and the column was allowed to re-equilibrate for 30 min. The flow rate was fixed at 50 μL/min, and 5 μL of each sample was injected. A Thermo Scientific Q Exactive Plus Orbitrap Mass Spectrometer (San Jose, CA, USA) was interfaced with the UHPLC system using a pneumatic-assisted heated electrospray ion source. Nitrogen was used as the sheath and auxiliary gases, which were set at 15 and 5 arbitrary units, respectively. The auxiliary gas was heated to 200°C. The heated ESI probe was set to 4000 V, and the ion transfer tube temperature was set to 300°C. Mass spectrometry (MS) detection was performed in positive ion mode operating in TOP-10 data dependent acquisition (DDA) mode. A DDA cycle entailed one MS^1^ survey scan (m/z 400-1500) acquired at 70,000 resolution (FWHM) and precursor ions meeting the user-defined criteria for charge state (i.e., z = 2, 3 or 4), monoisotopic precursor intensity (dynamic acquisition of MS2-based TOP-10 most intense ions with a minimum 1×104 intensity threshold) was selected for MS^2^. Precursor ions were isolated using the quadrupole (1.5 Da isolation width), activated by HCD (28 NCE) and fragment ions were detected in the Orbitrap at a resolution of 17,500 (FWHM). Data were processed using Thermo Proteome Discoverer (version 2.4) in conjunction with SEQUEST using default settings unless otherwise specified. The identification of peptides and proteins with SEQUEST was performed based on the reference proteome extracted from UniProt (*Mus musculus* taxon identifier 10090) as well as using a customized database including cytokines and chemokines protein sequences as FASTA sequences. Parameters were set as follows: MS^1^ tolerance of 10 ppm; MS^2^ mass tolerance of 0.02 Da for Orbitrap detection; enzyme specificity was set as trypsin with two missed cleavages allowed; carbamidomethylation of cysteine was set as a fixed modification; and oxidation of methionine was set as a variable modification. The minimum peptide length was set to six amino acids. Data sets were further analyzed with Percolator. Peptide-spectrum matches (PSMs) and protein identification were filtered at a 1% false discovery rate (FDR) threshold. For protein quantification and comparative analysis, we used the peak integration feature of Proteome Discoverer 2.4 software. For each identified protein, the average ion intensity of unique peptides was used for protein abundance.

#### 2.5 Bioinformatics

Abundance Ratio (log_2_): (experimental group)/(control group), Abundance Ratio p-value: (experimental group)/(control group) and Accession columns were extracted from the datasets generated by Proteome Discoverer. Volcano plots were generated using all identified and quantified proteins with both a log_2_ ratio and p-value. Proteins with a grouped abundance value of 0, without a p-value, or with a p-value > 0.05 were not included for further functional analysis. Gene Ontology analysis of genes (i.e., GO terms) was performed using Metascape [75], where significant proteins with a −log_10_ p-value ≥ 1.301 (p < 0.05) were used. The results from the upregulated and downregulated protein analyses were used to create bar plots in GraphPad Prism using the Metascape reports with the number of Differentially Expressed Proteins (DEPs) of GO process matches given by Metascape. Further pathway analysis was visualized with Cytoscape [55] using the GO, Kyoto Encyclopedia of Genes and Genomes (KEGG), REACTOME and WikiPathways pathway databases. Significant pathways for proteins were identified, comparing the ratio of target proteins identified in each pathway to the total number of proteins within the pathway. The statistical test used to determine the enrichment score for KEGG or REACTOME pathways was based on a right-sided hypergeometric distribution with multiple testing correction [4]. The clustered heat map was generated using R (https://cran.r-project.org/) with the package “pheatmap” (https://cran.r-project.org/web/packages/pheatmap/).

#### 2.6 BMCE Preparation and Administration

Bone marrow (BM) cells were harvested from both male and female 2 – 6 months old C57BL/6 mice. BM cells were flushed from the humeri, tibias and femurs with PBS and strained through a 40-μm filter. Erythrocytes were lysed with Ammonium-Chloride-Potassium (ACK) buffer, and the cells were washed again with 1x PBS. The cells were lysed with three cycles of freeze-thawing at −80°C and 37°C and centrifuged at 16 000 x g for 30 min at 4°C to remove insoluble materials. Protein concentration was assessed with a bicinchoninic acid (BCA) protein assay (Thermo Scientific), and subsequently diluted to the appropriate concentration with saline. BMCE (100 μL) contained 2 mg/mL protein was administered intravenousely to PSNL mice (n=5), twice a week, for 3 weeks. Another group of 5 PSNL mice received saline injections (used as the vehicle control).

#### 2.7 Statistical Analysis

Data are presented as mean ±SEM and analyzed using the GraphPad Prism software. The 2-tailed paired t test was used to analyze sham serum transfer behavior data, while the repeated-measures oneway analysis of variance (ANOVA) followed by the Dunnett post hoc test was used for PSNL serum transfer behavior data. Behavioral data for BMCE treatment was analyzed with a two-way ANOVA followed by the Sidak post hoc test. An ordinary one-way ANOVA with Tukey post hoc test was used to analyze behavioral data from sham, PSNL, and BMCE treated serum transfer. Differences were considered significant at p < 0.05.

## Results

### Peripheral nerve injury alters mouse serum proteome shortly after the insult

Although several previous clinical studies described the presence of systemic inflammation in chronic pain patients, they were limited to specific cytokine/chemokine targets, using ELISAs and multiplex immunoassays. To have a broader coverage and greater sensitivity to delineate changes taking place at the systemic level following nerve injury, we assessed serum proteome with label-free proteomics. Samples subject for analysis were obtained by pooling serum from 6 mice at 1 day after PSNL or sham surgery. Three panels, including, total serum proteins, immunoglobulin (Ig) clusters and cytokines/chemokines were examined. The volcano plot in **Fig. 1A** depicted differential abundances (PSNL vs sham), with the log_2_ ratio on the x-axis representing the fold change and the −log_10_ (p-value) on the y-axis depicting significance. A horizontal line represents the position for a p-value of 0.05, and the two vertical lines represent the positions of the 1.0 and −1.0 log2 ratios that correspond to two-fold increases/decreases coloured blue or red respectively. The coloured points, which pass these filters, are considered differentially expressed proteins (DEPs). At 1 day-post injury, 97 serum proteins were significantly altered, with 66 were upregulated and 31 were downregulated. Gene Ontology (GO) and Pathway enrichment analyses using Metascape were performed only on these significant DEPs. Summary connectivity maps were generated for enriched terms (**Fig. 1B-C**), where terms with a similarity >0.3 were connected by edges. In the first network plot (**Fig. 1B**), each node represents an enriched term and is colored by cluster ID, where nodes sharing the same cluster ID would be closer to each other. The second network plot (**Fig. 1C**) shows the same nodes but each one is colored by p-value, where the better represented pathways containing higher proportions of proteins involved in the pathway would have a more significant p-value. GO annotation indicated that the DEPs were mainly enriched in complement and coagulation cascades, complement cascade, negative regulation of proteolysis, inflammation, and hemostasis related pathways. The number of DEPs in each pathway was plotted in order from least to most significant p-values in **Fig. 1D**. Detailed description of > 2 fold-up (orange)- and down (green)-, and <2 fold (blue) significantly regulated enriched terms was summarized in supplementary table 1. The heatmap **in Fig. 2** illustrated changes in Ig clusters, in both variable (**Fig. 2A**) and constant (**Fig. 2B**) regions. Compared with serum in sham group, mainly Ig k light chains, a few λ light chains and Ig heavy chains were found to be upregulated in day 1-PSNL serum. It is interesting to notice that Ig J chain in the constant region, required for Ig M or Ig A secretion, is increased by nerve injury. The volcano plot in **Fig. 2C** depicted differential abundances of chemokines and cytokines in day-1 serum. At the acute phase post-PSNL, 13 cytokines and chemokines were significantly increased, where most of them have pro-inflammatory properties. Surprisingly, classic IL-1β, TNF-α and IL-6 were not among those significantly up-regulated cytokines. This result demonstrated that, a partial ligation on one of the sciatic nerve resulted quickly in a robust disturbance in serum homeostasis, with a strong upregulation of pro-inflammatory cytokines/chemokines, an increase of Ig clusters and regulation of around 100 serum proteins, mainly involved in inflammation-related pathways.

**Figure 1.**
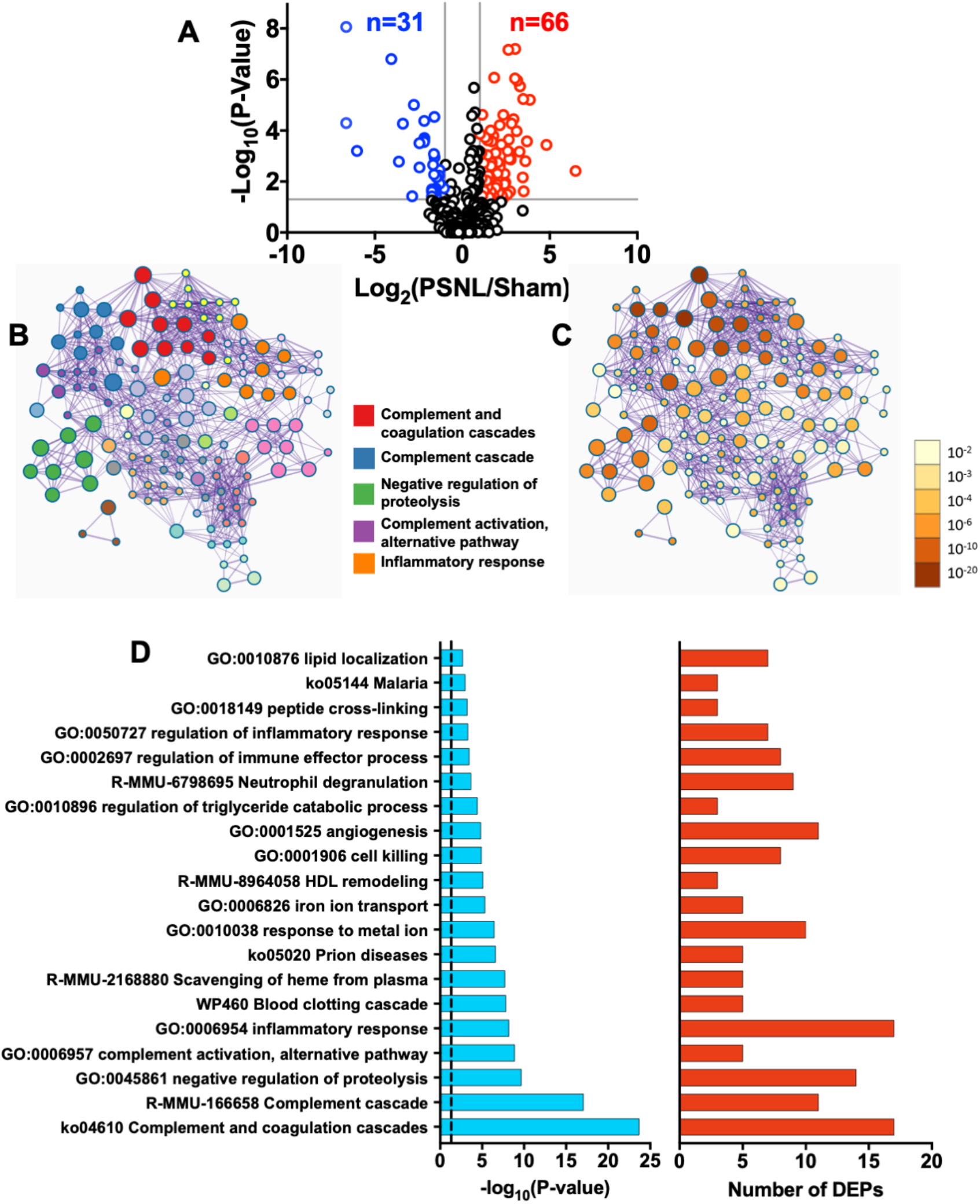
Mouse serum proteomics at 1-day post-PSNL. (A) Volcano Plot showing the differential abundances of total proteins in PSNL serum compared to sham serum. The x-axis represents the log2 ratio, with respect to Sham, and the y-axis represents the −log10 of the p-value. A horizontal line depicts the position of p-value at 0.05 and two vertical lines depict the position of 1.0 and −1.0 log2 ratios (corresponding to 2-fold increase/decrease, colored in red/blue respectively). (B-C) Network analysis of enriched pathways from Gene ontology (GO), Kyoto Encyclopedia of Genes and Genomes (KEGG), REACTOME, and WikiPathways, shown as a connectivity map of a subset of terms selected with the best p-values from each cluster. Analysis was performed with all significantly changing DEPs (upregulated and downregulated) in PSNL serum compared to Sham serum. Terms with a similarity >0.3 were connected by edges. The size of each node represents the number of DEPs represented within that enriched term. (B) Network plot where each node represents an enriched term and is colored by cluster ID, where nodes sharing the same cluster ID would be closer to each other. (C) The same network plot where each node is colored by p-value, where terms containing more genes would have a more significant p-value. (D) Log10 p-values and numbers of DEPs for each pathway cluster represented in the connectivity map. The vertical dashed line represents the position of p-value of 0.05. n = 6 animals/group. PSNL: partial sciatic nerve ligation.

**Figure 2.**
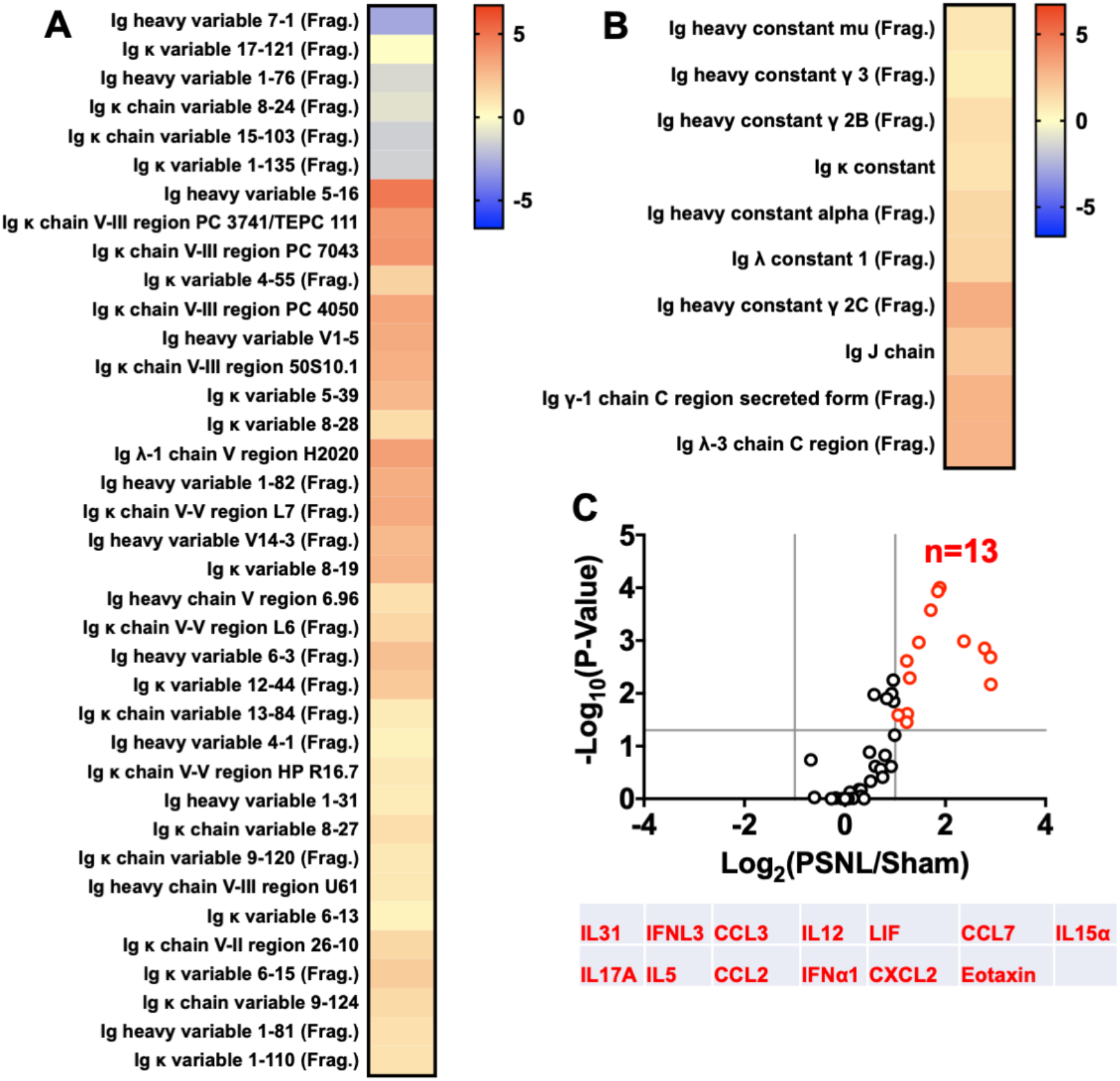
Mouse serum immunoglobulin, cytokine, and chemokine levels at 1-day post-PSNL. (A-B) Heatmap of log2 abundance ratios for immunoglobulin in PSNL and sham serum, separated by (A) variable region peptides and (B) constant region peptides. (C) Volcano plot showing the differential abundances of cytokines and chemokines in PSNL serum compared to sham serum. A list of all significant differentially expressed cytokines and chemokines is provided below the plot. n = 6 animals/group. PSNL: partial sciatic nerve ligation

### Nerve injury induced disturbance in mouse serum proteome persists for at least 1-month

To further understand the long term impact of nerve injury on systemic homeostasis, we performed another proteomic analysis on serum collected at one-month post-PSNL. The volcano plot (**Fig. 3A**) for total proteins showed 28 upregulated and 44 downregulated proteins, suggesting that although less abundant, 72 proteins still remained significantly altered at the late phase. Importantly, the most significant pathways involving enriched DEPs remained associated with inflammation, such as in complement activation and acute inflammatory response pathways, where DEPs in complement activation pathway were highly significant (**Fig. 3B-D**). Supplementary table 2 showed a detailed description of > 2 fold-up (orange)- and down (green)-, and <2 fold (blue) significantly regulated enriched terms. Again although less intense than those at acute phase, some slight up- and down-regulations were still detected in both variable (**Fig. 4A)** and constant (**Fig. 4B)** regions of Ig clusters, consisting primarily of Ig k light chains and Ig heavy chain variable regions. The volcano plot in **Fig. 4C** depicted 8 cytokines and chemokines were decreased and there included pro-inflammatory molecules and Th2 cells derived anti-inflammaotry cytokine IL-4 and IL-9. The analysis of late phase serum from PSNL mice clearly indicated that following a nerve injury, serum proteome profile at chronic stage is very different from that in acute phase. One month after the nerve injury, systemic inflammation appears less intense than that in acute phase, but it persists, and is mediated by different pathways.

**Figure 3.**
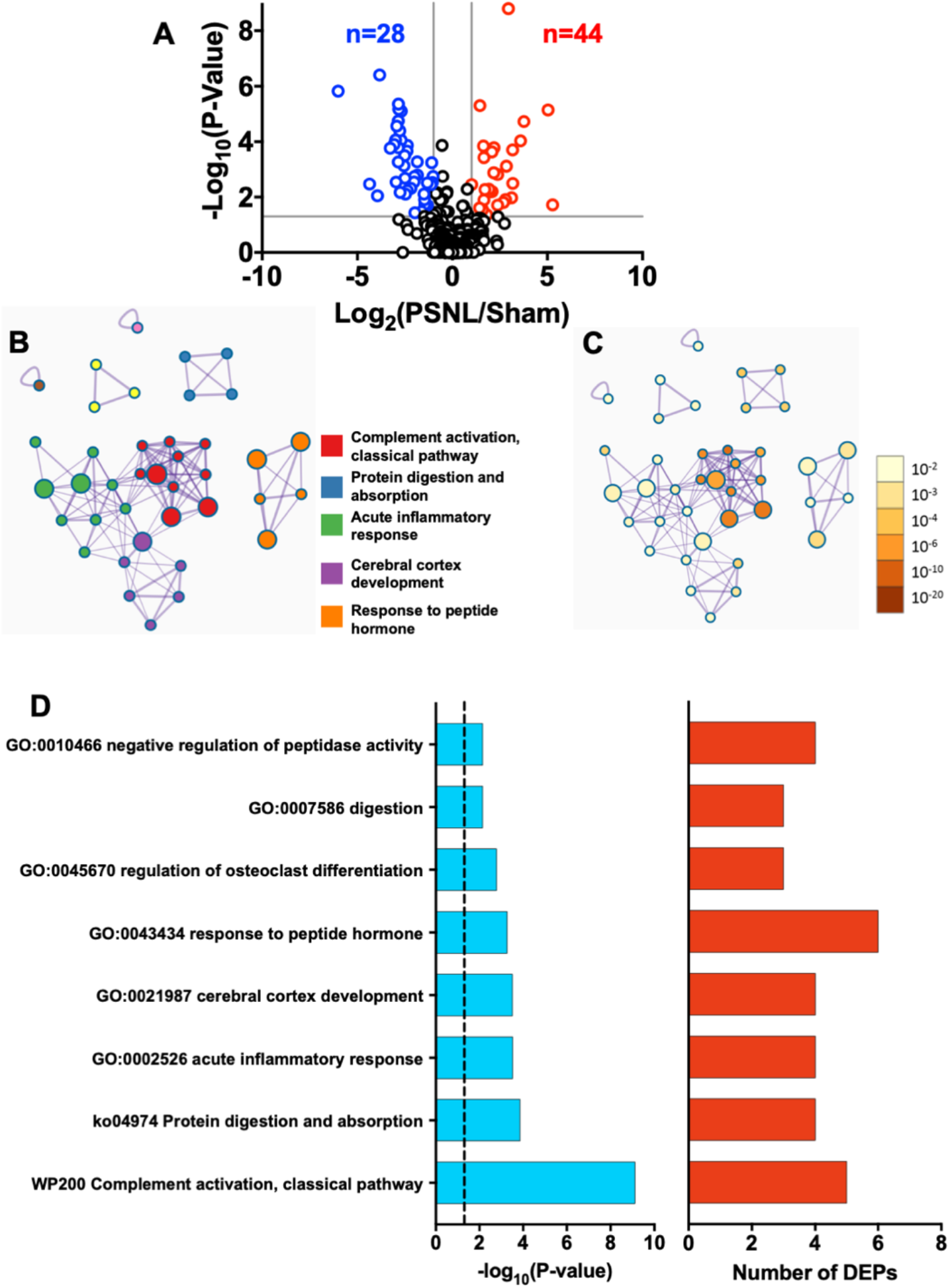
Mouse serum proteomics at 1-month post-PSNL. (A) Volcano Plot showing the differential abundances of cytokines and chemokines in PSNL serum compared to sham serum. (B-C) Network analysis of enriched pathways from Gene ontology (GO), Kyoto Encyclopedia of Genes and Genomes (KEGG), REACTOME, and WikiPathways, shown as a connectivity map of a subset of terms selected with the best p-values from each cluster. Analysis was performed with all significantly changing DEPs (upregulated and downregulated) in PSNL serum compared to Sham serum. The size of each node represents the number of DEPs represented within that enriched term. (B) Network plot where each node represents an enriched term and is colored by cluster ID, where nodes sharing the same cluster ID would be closer to each other. (C) The same network plot where each node is colored by p-value, where terms containing more genes would have a more significant p-value. (D) Log10 p-values and numbers of DEPs for each pathway cluster represented in the connectivity map. The vertical dashed line represents the position of p-value of 0.05. n = 6 animals/group. PSNL: partial sciatic nerve ligation

**Figure 4.**
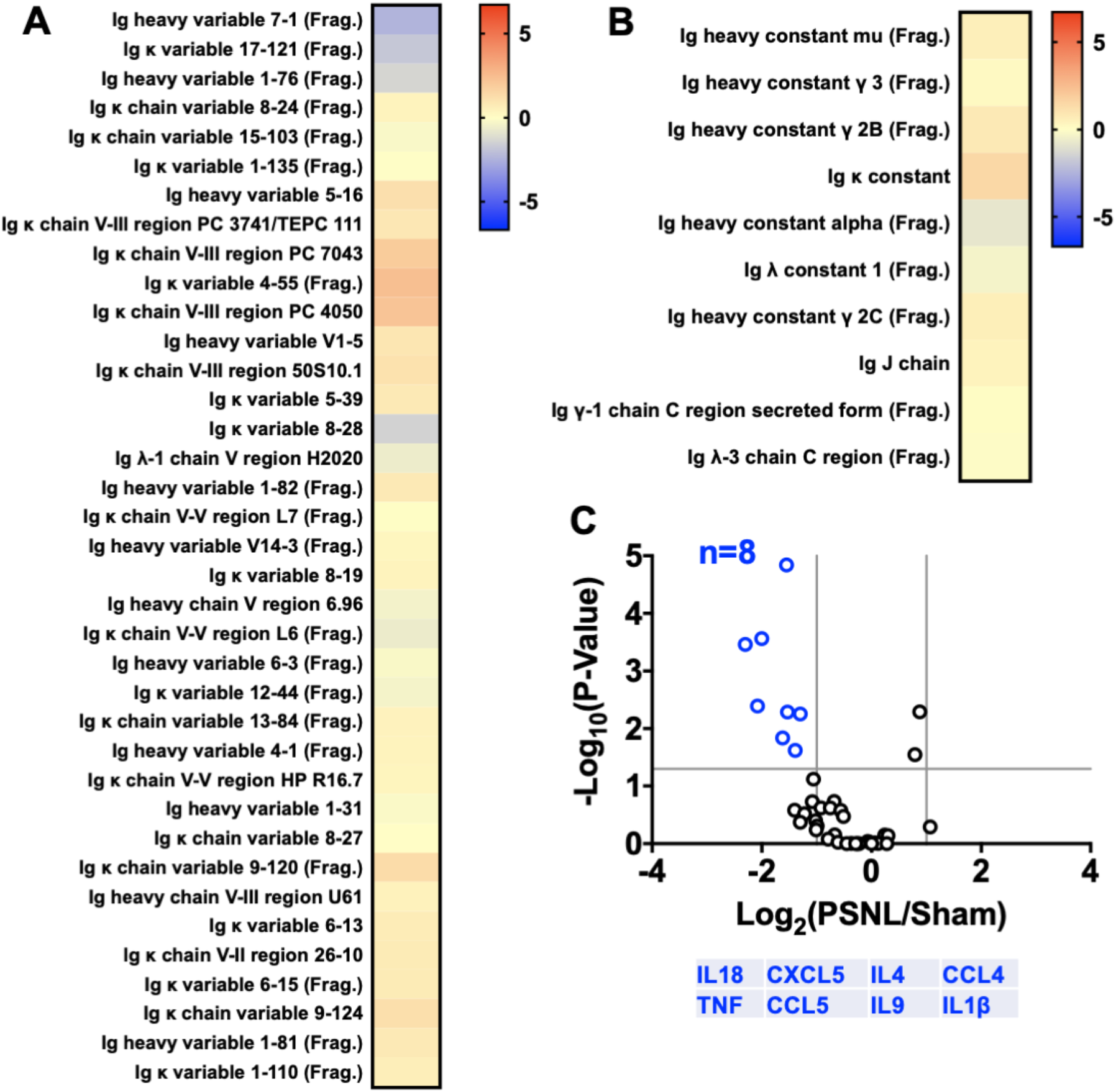
Mouse serum immunoglobulin, cytokine, and chemokine levels at 1-month post-PSNL. (A-B) Heatmap of log2 abundance ratios for immunoglobulin in PSNL and sham serum, separated by (A) variable region peptides and (B) constant region peptides. (C) Volcano plot showing the differential abundances of cytokines and chemokines in PSNL serum compared to sham serum. A list of all significant differentially expressed cytokines and chemokines is provided below the plot. n = 6 animals/group. PSNL: partial sciatic nerve ligation

### Serum from PSNL mice triggers painful response in naïve mice

Intrigued by the functional meaning of such changes in the serum of mice with nerve injury, we transferred serum from PSNL mice to naïve mice and monitored their pain behavior. By using von Frey and acetone tests respectively, our data showed that serum from sham-operated mice, either from 1-day or 1-month post-surgery, did not significantly affect mechanical thresholds and cold sensitivity in naïve mice (**Fig. 5A**). However, transferring PSNL serum (day 1-post injury) to naïve mice induced mechanical and cold allodynia, which lasted 3-4 days (**Fig. 5B**). Interestingly, it appears that serum from 1-month post-PSNL mice induced mechanical and cold allodynia that lasted slightly longer than that of acute serum, up 5-6 days (**Fig. 5C**). Furthermore, PSNL serum induced mechanical hypersensitivity was not restricted only to the paw but has also been detected in another area of the body, for example, in the orofacial area, this mechanical allodynia lasted for 4 days (**Fig. 5D**).

**Figure 5.**
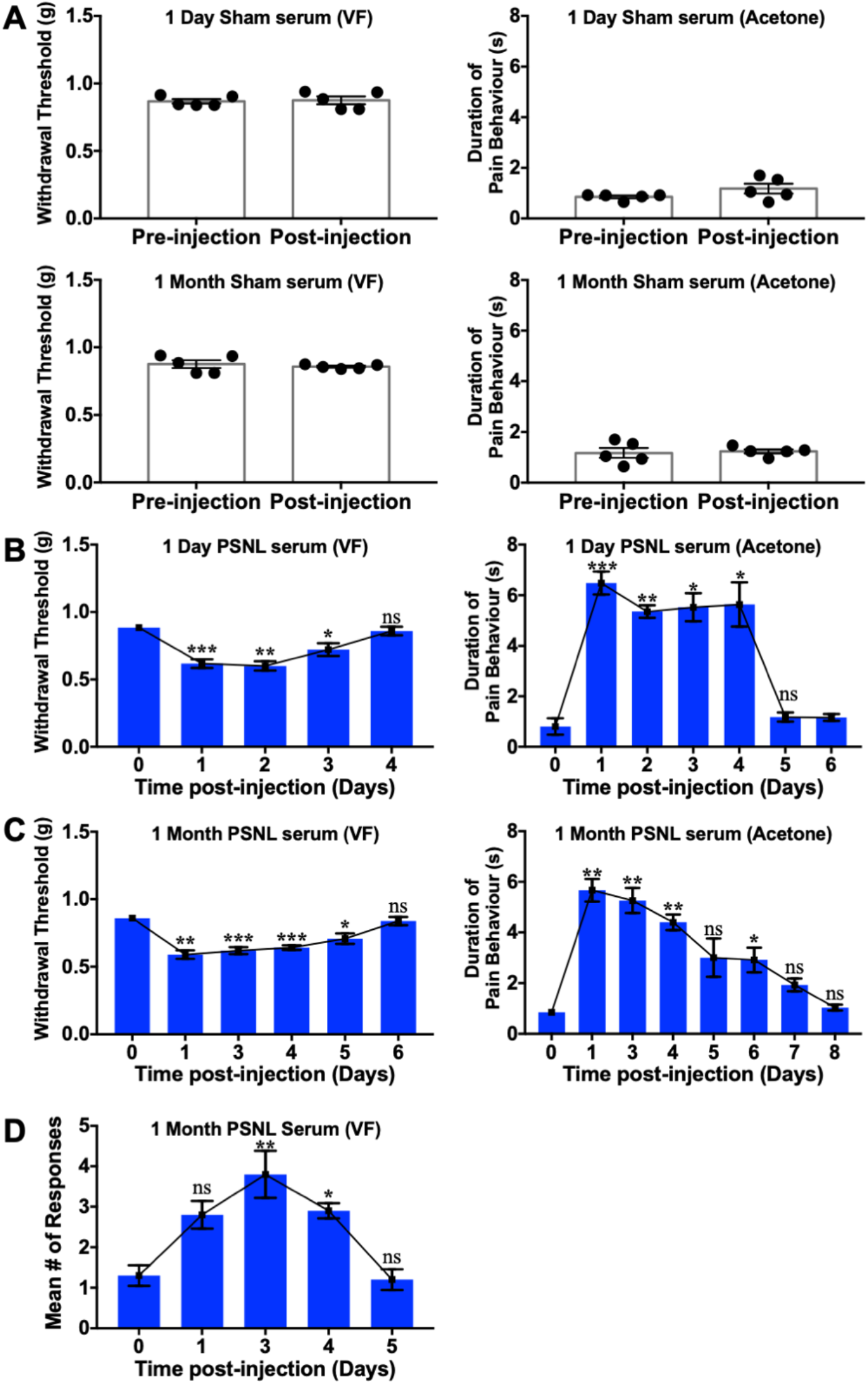
Serum from PSNL mice triggers painful response in naïve mice. (A) No significant changes in pain behavior were observed following i.v. administration of sham serum, tested at 1-day post-injection. Serum transferred from both (B) day 1 and (C) day 31 post-PSNL mice decreased mechanical threshold and increased cold sensitivity in naïve mice. Day 31 serum-triggered painful behavior lasts longer than that of day 1. (D) Serum from day 31 post-PSNL mice transferred to naïve mice increased mechanical hypersensitivity in the facial area. Data are mean ± SEM. n = 5 animals/group. *p < 0.05, **p < 0.01, ***p < 0.001, paired two-tailed Student’s t test (A) and one-way RM ANOVA with Dunnett’s post hoc test (B-D). PSNL: partial sciatic nerve ligation; BL: baseline.

### BMCE treatment significantly impacts mouse pain behavior

Given that there is a broad systemic change in the serum proteome following nerve injury and that this serum has pronociceptive properties, we wanted to assess whether a multifaceted systemic approach such as BMCE treatment could alleviate pain by dampening the nerve injury-induced serum proteomic disturbances. Twice a week intravenous administration of BMCE for 3 weeks significantly reduced mechanical (**Fig. 6A**) but not cold allodynia in PSNL mice (**Fig. 6B**). Furthermore, serum taken from PSNL mice treated with BMCE did not induce mechanical allodynia (**Fig. 6C**), but slight cold allodynia (**Fig. 6D**) when transferred to naïve mice, whereas serum from PSNL mice treated with saline did induce both mechanical and cold allodynia in naïve mice (**Fig. 6C-D**).

**Figure 6.**
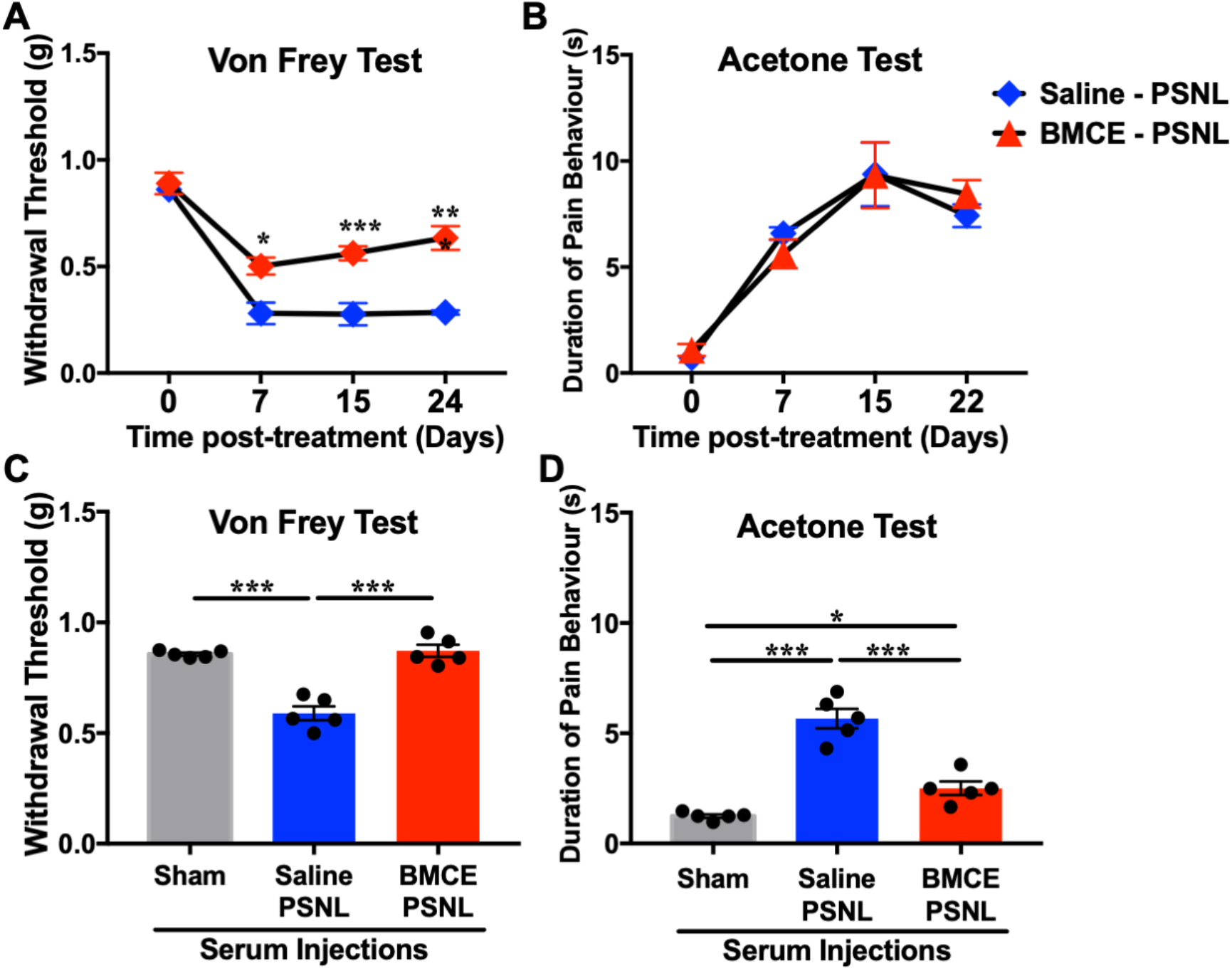
BMCE treatment reduces mechanical allodynia but not cold hypersensitivity. Behavioral analysis of BMCE treatment in PSNL mice with the (A) von Frey and (B) acetone tests. BMCE treatment 2 times per week starting on the day of nerve injury significantly reduced mechanical allodynia but did not impact cold sensitivity. Serum transferred from BMCE-PSNL mice (C) alleviated mechanical and (D) cold hypersensitivity induced by PSNL serum in naïve mice. Data are mean ± SEM. n = 5-10 animals/group. *p < 0.05, **p < 0.01, ***p < 0.001, two-way RM ANOVA with Sidak’s post hoc test (A-B) and one-way ordinary ANOVA with Sidak’s post hoc test (C-D). PSNL: partial sciatic nerve ligation; BL: baseline.

### BMCE treatment partially restores the homeostasis of proteomic profile in PSNL mice

To elucidate the systemic effects of BMCE treatment on nerve injury and to understand underlying mechanisms of its effects in alleviating pain in PSNL mice, serum proteome of BMCE-treated PSNL mice was analyzed. To compare with acute (**Fig. 1A**) and chronic phase (**Fig. 3A**), there were less proteins remain altered, totalling 57 proteins, where 37 proteins were upregulated and 17 were downregulated (**Fig. 7A**). GO annotation indicated that the most significant pathways involving enriched DEPs with BMCE treatment were in acute-phase response, complement and coagulation cascades, negative regulation of peptidase activity, and gland development pathways (**Fig. 7B**). In general, to compare with acute or chronic phase PSNL serum, not only less number of DEPs were involved in the different pathways, depicted as a less significant p-value (**Fig. 7C**), and but most of the pathways, except the complement pathway with few DEPs, are not related to inflammation (**Fig. 7B, D**). Detailed description of > 2 fold-up (orange)- and down (green)-, and <2 fold (blue) significantly regulated enriched terms was summarized in supplemantery table 3. Some Ig clusters, involving Ig heavy and Ig κ variables, were found increased in BMCE/PSNL serum, mainly in the variable regions (**Fig. 8A, B**). Only 3 cytokines were found significanly up-regulated with BMCE treatment, which is dominated by the increase of IL-27 (**Fig. 8C**), and it has potential in inducing the production of IL-10. The cytokine/chemokine heatmap (**Fig. 8D**) revealed that BMCE treatment tends to re-establish inflammatory balance, where strong up- or down-regulation was no longer detected in BMCE/PSNL serum. Altogether it suggests that 3 week-BMCE treatment inhibited systemic inflammation and partially restored serum homeostasis disturbed by nerve injury.

**Figure 7.**
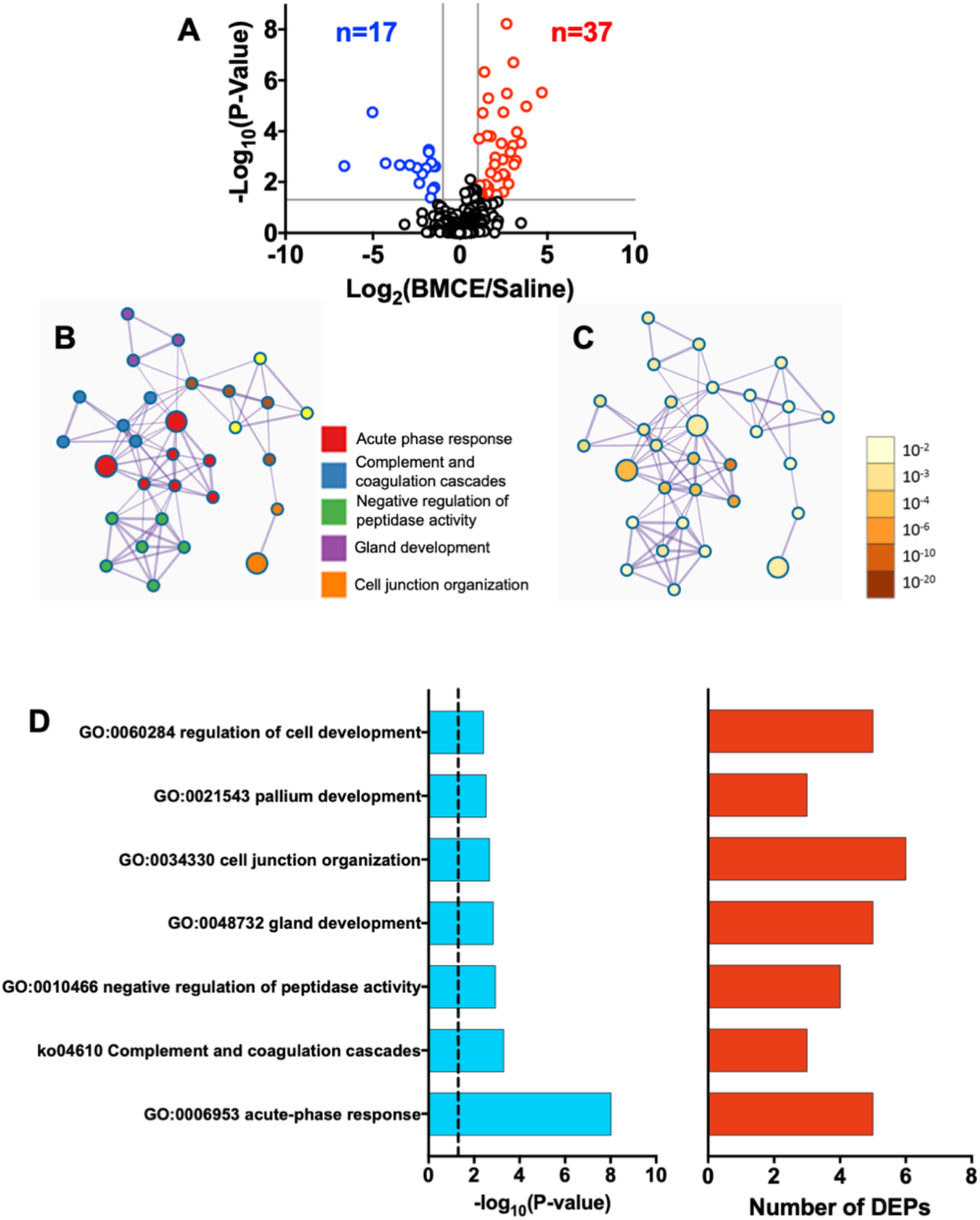
BMCE treatment partially restores the serum proteomic profile in PSNL mice. (A) Volcano Plots showing the differential abundances of total proteins in BMCE treated PSNL serum compared to saline treated PSNL serum. (B-C) Network analysis of enriched pathways from Gene ontology (GO), Kyoto Encyclopedia of Genes and Genomes (KEGG), REACTOME, and WikiPathways, shown as a connectivity map of a subset of terms selected with the best p-values from each cluster. Analysis was performed with all significantly changing DEPs (upregulated and downregulated) in BMCE treated PSNL serum compared to saline treated PSNL serum. The size of each node represents the number of DEPs represented within that enriched term. (B) Network plot where each node represents an enriched term and is colored by cluster ID, where nodes sharing the same cluster ID would be closer to each other. (C) The same network plot where each node is colored by p-value, where terms containing more genes would have a more significant p-value. (D) Log10 p-values and numbers of DEPs for each pathway cluster represented in the connectivity map. The vertical dashed line represents the position of p-value of 0.05. n = 5 animals/group. PSNL: partial sciatic nerve ligation.

**Figure 8.**
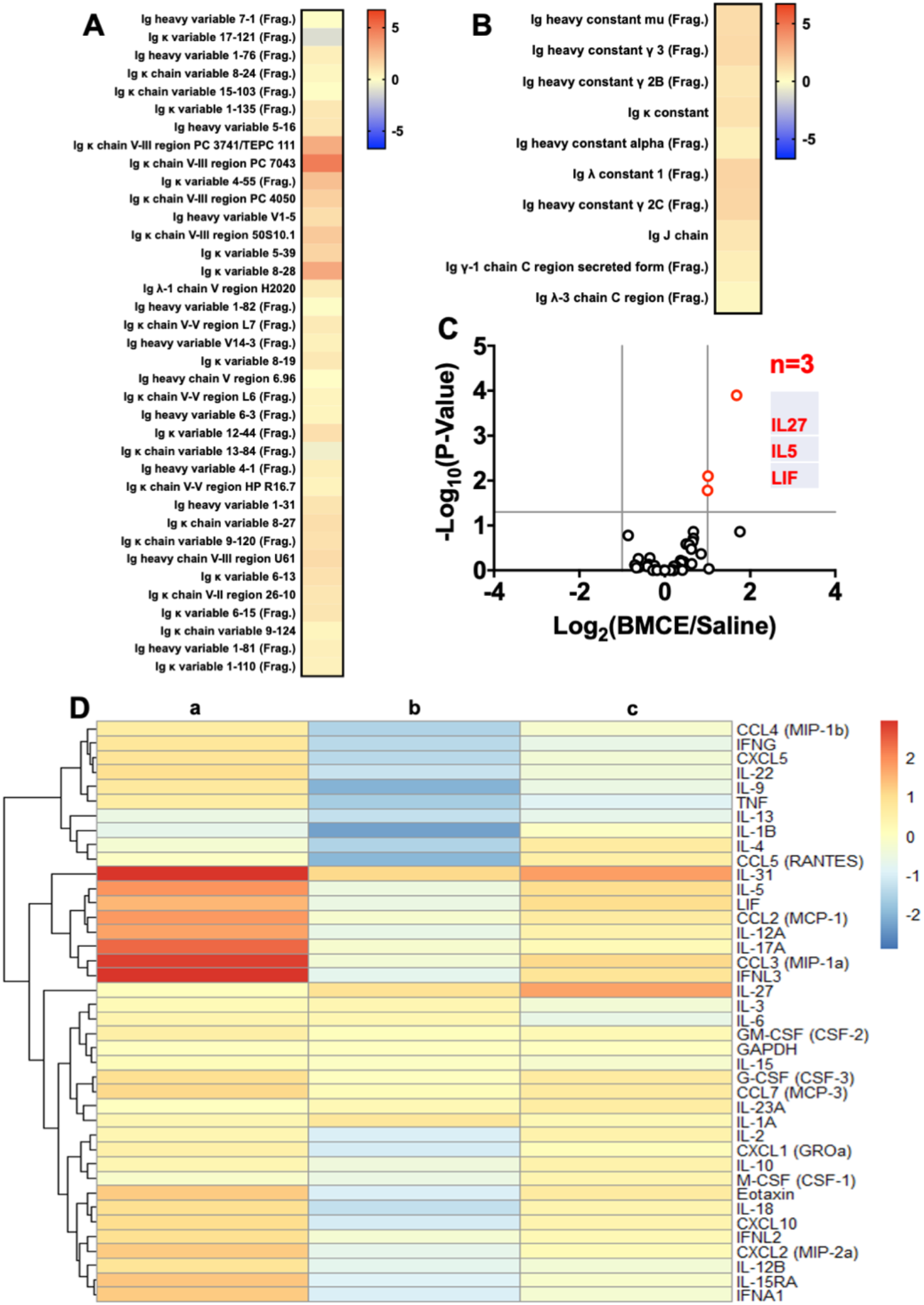
BMCE treatment partially restores serum immunoglobulin, cytokine, and chemokine levels in PSNL mice. (A-B) Heatmap of log2 abundance ratios for immunoglobulin in BMCE treated PSNL serum and saline treated PSNL serum, separated by (A) variable region peptides and (B) constant region peptides. (C) Volcano plot showing the differential abundances of cytokines and chemokines in BMCE treated PSNL serum compared to saline treated PSNL serum. A list of all significant differentially expressed cytokines and chemokines is provided below the plot. (D) Clustered heatmap drawn using R package “pheatmap” of log2 abundance ratios for chemokines and cytokines in (a) PSNL/Sham serum at 1-day post-PSNL, (b) PSNL/Sham serum at 1-month post-PSNL, and (c) BMCE treated PSNL serum/Saline treated PSNL serum at 3 weeks post-PSNL. n = 5-6 animals/group. PSNL: partial sciatic nerve ligation.

## Discussion

By using a high throughput, unbiased and semi-quantitative approach of serum proteome profiling, we demonstrated that nerve injury resulted in a long lasting systemic disturbance with a systemic inflammation predominance. We also addressed the potential of systemic factors contributing to neuropathic pain. Transferring serum from nerve injured mice into naïve mice also transferred mechanical and cold hypersensitivities. While BMCE partially re-established systemic homeostasis in nerve injured mice, it alleviated neuropathic mechanical hypersensitivity and serum from BMCE-treated-nerve-injured mice corrected the painful effect of nerve injured serum. Altogether, our data strongly suggest the involvement of systemic modulation in the pathophysiology of chronic neuropathic pain.

The potential of systemic contribution in chronic pain has drawn attention in recent years [74]. Some inflammatory mediators, mainly cytokines/chemokines, have been repeatedly detected in the blood of patients suffering from chronic pain with different pathologies, including injury associated peripheral neuropathy [28; 37; 61]; complex regional pain syndrome (CRPS) [60]; diabetic neuropathy [11]; and also in chronic widespread pain (CWP) patients [20; 62; 65]. While most of the above mentioned studies analysing mRNA expression or protein levels of few selected targets in blood samples, some studies [2; 29; 34] expanded the screening of inflammatory molecules with multiplex analysis up to 92 targets where they not only detected significant differences in multiple inflammatory biomarkers, but also observed that among patients whose pain and other clinical relevant symptoms improve, most of inflammation related proteins tend to be normalized [29]. Most recently, a series of studies using high-throughput serum/plasma proteomics further revealed a dysregulation of circulating proteome in CWP, especially fibromyalgia (FM) patients, where the most regulated proteins are part of the immune system or known to interact with the immune system [21; 22; 27; 64–67]. The data further support a significant correlation between serum/plasma proteome profile and clinical parameters in FM patients, such as mechanical and thermal pain thresholds, pain intensity and psychological distress [21; 22; 64; 65]. Thus, accumulating evidence suggests that chronic pain patients may live with systemic chronic inflammation, which could have a role in the genesis of long lasting painful behhaviors.

In the current animal study, by using unbiased proteomic analysis, we demonstrated that a partial ligation on one of the sciatic nerves led to a long lasting disturbance in serum proteome. A robust systemic inflammatory reaction occured one day after the nerve injury. Among others, a significant increase of at least 13 proinflammatory cytokines/chemokines, an increase of multiple variables in immunoglobin clusters and a sturdy disturbance in complement pathways were detected. One month after the nerve ligation, while a down-regulated cytokine/chemokine profile predominated and the alteration in Ig clusters appeared less important, the changes in inflammation related pathways remained highly significant, especially the DEPs involved in the complement activation pathways. Thus, we confirmed that nerve injury is associated with a systemic chronic inflammation, with a distinct inflammatory signature at the acute and chronic phase. Our serum transfer experiments validated the functional significance of these systemic factors in the serum, as they indeed induced mechanical and cold hypersensitivity in naïve mice. Different temporal profile of pain induced by 1-day or 1-month-PSNL serum in naïve mice can actually be explained by the different compostions in these sera. We may assume that in addition to many other altered inflammation related proteins, cytokines/chemokines are important players in 1-day-serum induced hypersensitivity, while proteins in complement pathways are essential in 1-month-serum induced enhanced pain behavior.

Although nerve injury is one of the major reasons often causing debilitating neuropathic pain, our understanding of underlying mechanisms still remains poor. It can involve different types of cells within the nervous system, it can also interact with other systems, especially the immune system, leading to peripheral and/or central sensitization [31; 32]. Inflammatory systemic factors that we detected in the serum of nerve injured mice could be derived from active release from the circulating immune cells or/and leakage from distressed or damaged cells of the surrounding organs/tissues. As a complex pathological process, neuropathic pain rarely results from any one single serum protein acting alone rather through highly interacting newtworks of multiplayers driven by the dynamics of the local environment. To this end, to further confirm the contribution of systemic inflammation to neuropathic pain, we used a broad targeting strategy to treat nerve injured mice with BMCE which has established anti-inflammatory and regenerative properties via its paracrine effects [57; 59]. It has been noticed that while our proteomic analysis clearly indicated that BMCE was able to, at least partially, re-establish inflammatory balance and restore systemic serum homeostasis in PSNL mice, and it alleviated nerve injury triggered mechanical allodynia. However, it was not effective in reducing cold hypersensitivity. This is in coincidence with our serum transfer experiment where BMCE treatment completely erased the effect of PSNL serum in inducing mechanical allodynia in naïve mice, but it still had some effects in triggering cold allodynia. This distinctive response of BMCE in different pain modalities is indeed inline with a recent finding where the authors claimed that immune system activation contributes to tactile allodynia but minimally to cold allodynia [6].

While we confirmed the role of systemic inflammation in neuropathic pain, it remains to be addressed how systemic factors in the bloodstream contribute to increasing neuronal sensitization. Needless to argue that as inflammatory mediators, cytokines/chemokines can not only directly excite neurons, but also influence the function of non-neuronal cells in the nervous system, including the integrity of the neural barriers and PNS/CNS glial cell activation [19; 48; 49; 53]. The contribution of cytokines/chemokines in chronic pain has been well-established with a large amount of pre-clinical and clinical evidence [15; 19; 29; 30; 53; 73]. In recent years, there is a growing appreciation that autoimmunity could be one of the mechanisms for chronic pain [36]. In randomized controlled trials for rheumatoid arthritis, infusions of rituximab to deplete B cells significantly improved patients’pain experience [13]. Depletion of B cells also reduced the severity of mechanical allodynia and vascular damage in an animal model of CRPS [42]. Transferring IgG from FM patients produced FM-like symptoms in mice, including mechanical and cold hypersensisitivities, reduced physical activities and paw grip strength [25]. Increases in several Ig variable regions have been detected in our study in the serum of both day-1 and day-30 PSNL mice. While unlikely this is an antigen specific adaptive activation of B cells at day-1 post-injury, it may reflect an activation of innate-like B cells (ILBs) triggered by DAMPs through TLR pathway. As unconventional B cells, ILBs have innate sensing and responsing properties and they can produce natural IgM to protect the host, for example, to promote the engulfment of tissue debris/apoptotic cells, before an adaptive immune response is elicited [14; 50]. Such hypothesis may be supported by the fact that we also detected an increase of Ig J chain in the constant region, which is required for the IgM pentameric structure [8]. However, the possibility of nerve tissue antigen-specific autoimmune response would not be excluded as we also found an increase of Ig variables at one-month post-injury, which may be relevant to an activation of conventional B cells and plasma cells, leading to the production of autoantibodies. Although with the current proteomic analysis, it is difficult to conclude whether antibodies are directly involved in neuropathic pain, anyhow it suggests the presence of B cell-mediated humoral immune response in nerve injured mice. The complement system has been implicated in different chronic pain conditions [69]. Various complement proteins were found elevated at the site of nerve injury, or in the spinal cord [41; 46]; and using either genetic or pharmacological approaches to block the activation reduced pathological pain [26; 43]. Complement factors, C3 and C5 [17; 24; 26; 40], were also found elevated in the plasma and cerebrospinal fluid of neuropathic pain patients and relevant animal models. Likewise, our proteomic anlysis detected a strong activation of complement pathways in the serum of nerve injured mice, both at acute and chronic phase, which was inhibited by BMCE and mice pain experience was improved in parallel.

## Conclusion

It is worth emphasizing that recent remarkable advances in protein profiling technologies (e.g., protein separation, mass spectrometry, and bioinformatics) offer great promise in pain research. Serum and plasma proteomics allow for monitoring the changes of functionally relevant proteins, and enables the interrogation of associated protein networks with their predicted activities [23; 45; 70]. In addition to providing clear evidence that nerve injury is assocated with a long term systemic disturbance, which has a role in the generation of chronic pain, the current findings also implicate multiple inflammation related players in pain pathology, including cytokines/chemokines, autoantibodies and complement factors. These molecules may act distinctively or synergetically, at different stages of the pathology, leading to chronic pain.

## Author contributions

WBSZ, XQS and YNL performed experiments; ST, FB and JZ conceived the study, and WBSZ, ST, FB and JZ wrote the manuscript

## Acknowledgements

This work was supported by funding from the Canadian Institutes for Health Research (CIHR) PJT-155929 to JZ, from the National Sciences and Engineering Research Council of Canada (NSERC) RGPIN-2020-05228 to FB and Fonds de recherche Quebec Sante (FRQS) JZ and FB. Proteomic equipment was funded by the Canadian Foundation for Innovation (CFI) and the Fonds de Recherche du Québec (FRQ) project #36706 to FB. The authors acknowledge technical assistance of Oladayo Oladiran and JingWen Meng.

