## Supplemental Table 1-3 for "Unbiased proteomic analysis detects painful systemic inflammatory profile in the serum of nerve injured mice"

**Supplementary table 1: Selected pathways for PSNL vs Sham serum 1-day post-surgery.** Pathways: (1) ko04610 Complement and coagulation cascades. (2) R-MMU-166658 Complement cascade. (3) GO:0045861 negative regulation of proteolysis. (4) GO:0006957 complement activation, alternative pathway. (5) GO:0006954 inflammatory response. (6) WP460 Blood clotting cascade. (7) R-MMU-2168880 Scavenging of heme from plasma. (8) ko05020 Prion diseases. (9) GO:0010038 response to metal ion. (10) GO:0006826 iron ion transport. (11) R-MMU-8964058 HDL remodeling. (12) GO:0001906 cell killing. (13) GO:0001525 angiogenesis. (14) GO:0010896 regulation of triglyceride catabolic process. (15) R-MMU-6798695 Neutrophil degranulation. Orange: > 2 fold significantly up-regulated enriched terms; Green: > 2 fold significantly down-regulated enriched terms; Blue <2 fold significantly up- and down-regulated enriched terms.

| Accession | Gene Name | Log2(PSNL /Sham) | P-Value | Description | Pathway(s) |
|---|---|---|---|---|---|
| P97481 | Epas1 | 3.61 | 0.001567618 | Endothelial PAS domain-containing protein 1 | 13 |
| E9Q861 | C2cd4a | 2.68 | 0.001247673 | C2 calcium-dependent domain-containing 4A | 5 |
| P28798 | Grn | 2.57 | 0.031769842 | Progranulin | 5, 13, 15 |
| P08071 | Ltf | 2.41 | 0.012724263 | Lactotransferrin | 3, 10, 12, 15 |
| P01592 | Igj; Jchain | 2.14 | 6.18188E-05 | Immunoglobulin J chain | 7 |
| Q05020 | Apoc2 | 2.14 | 0.002149921 | Apolipoprotein C-II | 11, 14 |
| P01887 | B2m | 1.9 | 0.000743563 | Beta-2-microglobulin | 9, 10, 12, 15 |
| Q8VCS0 | Pglyrp2 | 1.82 | 8.49901E-07 | N-acetylmuramoyl-L-alanine amidase | 5 |
| P51910 | Apod | 1.78 | 0.000162343 | Apolipoprotein D | 5 |
| Q8CG16 | C1ra | 1.69 | 0.000263318 | Complement C1r-A subcomponent | 1, 2 |
| O88947 | F10 | 1.64 | 0.025318017 | Coagulation factor X | 1, 6 |
| O88844 | Idh1 | 1.64 | 0.002402405 | Isocitrate dehydrogenase [NADP] cytoplasmic | 15 |
| Q02105 | C1qc | 1.59 | 0.007054343 | Complement C1q subcomponent subunit C | 1, 2, 8 |
| Q64726 | Azgp1 | 1.48 | 0.005874154 | Zinc-alpha-2-glycoprotein | 12 |
| A8DUK4 | Hbb-bs; Hbb-bt | 1.44 | 0.000912121 | Beta-globin | 7, 15 |
| P97298 | Serpinf1 | 1.31 | 0.000186498 | Pigment epithelium-derived factor | 3, 5, 13, |
| Q8VCU2 | Gpld1 | 1.14 | 2.3857E-05 | Glycosyl-phosphatidylinositol-specific phospholipase D | 9, 13, 14 |
| O09164 | Sod3 | 1.13 | 0.036545473 | Extracellular superoxide dismutase [Cu-Zn] | 9 |
| P05366 | Saa1 | 1.11 | 0.00761186 | Serum amyloid A-1 protein | 5 |
| Q61646 | Hp | 1.01 | 0.000134426 | Haptoglobin | 5, 7, 15 |
| P06683 | C9 | 0.98 | 0.003942248 | Complement component C9 | 1, 2, 4, 8, 12 |
| Q80YQ1 | Thbs1 | 0.96 | 0.000629717 | Thrombospondin-1 | 3, 5, 9, 13 |
| Q61247 | Serpinf2 | 0.93 | 0.010316519 | Alpha-2-antiplasmin | 1, 3, 5, 6 |
| F8WI14 | Ecm1 | 0.9 | 0.01340782 | Extracellular matrix protein 1 | 3, 5, 13 |
| Q03734 | Serpina3m | 0.9 | 0.005461573 | Serine protease inhibitor A3M | 3 |
| Q01339 | Apoh | 0.87 | 0.000719371 | Beta-2-glycoprotein 1 | 13, 14 |
| P16301 | Lcat | 0.85 | 0.020438894 | Phosphatidylcholine-sterol acyltransferase | 9, 11 |
| Q91X72 | Hpx | 0.84 | 8.33755E-05 | Hemopexin | 7, 10, |
| P03953 | Cfd | 0.83 | 0.009743108 | Complement factor D | 1, 2, 4, 15 |

| | | | | | |
|---|---|---|---|---|---|
| Q91X70 | C6 | 0.77 | 0.011839402 | Complement component 6 | 1, 8, 13 |
| P33587 | Proc | 0.75 | 0.00504131 | Vitamin K-dependent protein C | 1, 5 |
| Q9DBD0 | 1300017J02Rik | 0.74 | 0.00140022 | Inhibitor of carbonic anhydrase | 10 |
| P06909 | Cfh | 0.72 | 1.92348E-05 | Complement factor H | 1, 2, 4, 5, 12, 13 |
| O08677 | Kng1 | 0.71 | 0.022039689 | Kininogen-1 | 1, 3, 5, 12 |
| P28665 | Mug1 | 0.62 | 0.029207121 | Murinoglobulin-1 | 3 |
| Q01279 | Egfr | 0.61 | 0.000607702 | Epidermal growth factor receptor | 9 |
| Q61129 | Cfi | 0.59 | 0.002293686 | Complement factor I | 1, 2 |
| P01027 | C3 | 0.55 | 2.61882E-05 | Complement C3 | 1, 2, 4, 5, 12, 13, 15 |
| P19221 | F2 | 0.55 | 0.044735022 | Prothrombin | 1, 2, 3, 5, 6, 12 |
| Q9QXC1 | Fetub | 0.53 | 0.024444135 | Fetuin-B | 3 |
| A0A087WSN6 | Fn1 | 0.5 | 0.00076459 | Fibronectin | 5, 9, 13 |
| P29699 | Ahsg | 0.5 | 0.008371928 | Alpha-2-HS-glycoprotein | 3, 5, 15 |
| Q8K182 | C8a | 0.44 | 0.000215451 | Complement component C8 alpha chain | 1, 2, 4, 8 |
| Q07968 | F13b | 0.41 | 0.001393594 | Coagulation factor XIII B chain | 1, 6 |
| Q921I1 | Trf | 0.24 | 0.043358889 | Serotransferrin | 9, 10 |
| P07724 | Alb | -0.2 | 0.002953879 | Albumin | 7, 11 |
| Q9JJN5 | Cpn1 | -0.65 | 0.012535509 | Carboxypeptidase N catalytic chain | 2 |
| A0A0R4J0J1 | Cpne9 | -1.3 | 0.011909419 | Copine-9 | 9 |
| P70375 | F7 | -1.48 | 0.006783577 | Coagulation factor VII | 1, 6 |
| Q9JHH6 | Cpb2 | -1.74 | 0.037836572 | Carboxypeptidase B2 | 1, 2, 3 |
| P20029 | Hspa5 | -2.46 | 0.000310233 | Endoplasmic reticulum chaperone BiP | 8 |
| Q3UEL9 | Serpina7 | -3.4 | 5.34868E-05 | Thyroxine-binding globulin | 3 |

**Supplementary table 2: Selected pathways for PSNL vs Sham serum 1-month post-surgery.** Pathways: (1) WP200 Complement activation, classical pathway. (2) ko04974 Protein digestion and absorption. (3) GO:0002526 acute inflammatory response. (4) GO:0021987 cerebral cortex development. (5) GO:0043434 response to peptide hormone. (6) GO:0045670 regulation of osteoclast differentiation. (7) GO:0007586 digestion. (8) GO:0010466 negative regulation of peptidase activity. Orange: > 2 fold significantly up-regulated enriched terms; Green: > 2 fold significantly down-regulated enriched terms; Blue <2 fold significantly up- and down-regulated enriched terms.

| Accession # | Gene Name | Log2(PSNL/Sham) | P-Value | Description | Pathway(s) |
|---|---|---|---|---|---|
| P14106 | C1qb | 3.59 | 9.25942E-05 | Complement C1q subcomponent subunit B | 1 |
| Q8JZR2 | Crk | 3.11 | 0.010593704 | Adapter molecule crk | 4, 5 |
| Q61646 | Hp | 2.95 | 1.58738E-09 | Haptoglobin | 3 |
| Q04690 | Nf1 | 2.8 | 0.012385029 | Neurofibromin | 4, 6 |
| Q99N42 | Tymp | 2.09 | 0.006307418 | Thymidine phosphorylase | 7 |
| P15919 | Rag1 | 1.71 | 0.005367547 | V(D)J recombination-activating protein 1 | 8 |
| P07361 | Orm2 | 1.66 | 0.000375078 | Alpha-1-acid glycoprotein 2 | 3 |
| A2A9A2 | Dmrta2 | 1.63 | 0.000142638 | Doublesex- and mab-3-related transcription factor A2 | 4 |
| P01027 | C3 | -0.3 | 0.007218299 | Complement C3 | 1, 3 |
| P28665 | Mug1 | -0.51 | 0.011923339 | Murinoglobulin-1 | 8 |
| P06684 | Hc | -0.64 | 0.01292701 | Complement C5 | 1 |
| Q8BH35 | C8b | -0.64 | 0.041632049 | Complement component C8 beta chain | 1 |
| Q00897 | Serpina1d | -0.82 | 0.047953177 | Alpha-1-antitrypsin 1-4 | 5, 8 |
| Q06770 | Serpina6 | -0.88 | 0.007447719 | Corticosteroid-binding globulin | 8 |
| P00920 | Car2 | -1.33 | 0.019827465 | Carbonic anhydrase 2 | 5, 6 |
| Q00724 | Rbp4 | -1.33 | 0.019409515 | Retinol-binding protein 4 | 5, 7 |
| Q9D5V5 | Cul5 | -1.57 | 0.041928115 | Cullin-5 | 4 |
| Q63ZW6 | Col4a5 | -1.62 | 0.032963575 | Collagen type IV alpha 5 chain | 2 |
| Q91Y97 | Aldob | -1.85 | 0.000528666 | Fructose-bisphosphate aldolase B | 5 |
| E0CX52 | Clec2i | -2.02 | 0.002818325 | C-type lectin domain family 2 member I | 6 |
| Q8K1H9 | Obp2a | -2.55 | 0.000722974 | Odorant-binding protein 2a | 5 |
| E9Q861 | C2cd4a | -2.84 | 0.000164052 | C2 calcium-dependent domain-containing 4A | 3 |
| Q792Z1 | Try10 | -2.85 | 0.000535411 | Trypsin 10 | 2 |
| Q9CR35 | Ctrb1 | -3.94 | 0.0089315 | Chymotrypsinogen B | 2, 7 |
| Q9QUK9 | Try5 | -6.01 | 1.49421E-06 | TESP4 | 2 |

**Supplementary table 3: Selected pathways for BMCE vs Vehicle serum 3-week post-surgery.** Pathways: (1) GO:0006953 acute-phase response. (2) ko04610 Complement and coagulation cascades. (3) GO:0010466 negative regulation of peptidase activity. (4) GO:0048732 gland development. (5) GO:0060284 regulation of cell development. (6) GO:0034330 cell junction organization. Orange: > 2 fold significantly up-regulated enriched terms; Green: > 2 fold significantly down-regulated enriched terms; Blue <2 fold significantly up- and down-regulated enriched terms.

| **Accession** | **Gene Name** | **Log2(PSNL/ Sham)** | **P-Value** | **Description** | **Pathway(s)** |
|---|---|---|---|---|---|
| A2A9A2 | Dmrta2 | 3.05 | 1.98195E-07 | Doublesex- and mab-3-related transcription factor A2 | 6, 7 |
| P14106 | C1qb | 2.88 | 0.000667646 | Complement C1q subcomponent subunit B | 2, 5 |
| Q61646 | Hp | 2.66 | 5.99523E-09 | Haptoglobin | 1, 4 |
| P07361 | Orm2 | 2.47 | 1.79626E-05 | Alpha-1-acid glycoprotein 2 | 1 |
| Q6TA13 | Kif1a | 2.47 | 0.004770139 | Kinesin-like protein KIF1A | 5 |
| P15919 | Rag1 | 2.44 | 0.001288488 | V(D)J recombination-activating protein 1 | 3, 4 |
| Q04690 | Nf1 | 2.1 | 0.031352579 | Neurofibromin | 4, 6, 7 |
| Q9QY01 | Ulk2 | 2.1 | 0.006146829 | Serine/threonine-protein kinase ULK2 | 7 |
| P05366 | Saa1 | 1.75 | 0.000160267 | Serum amyloid A-1 protein | 1 |
| A0A0R4J0X5 | Serpina1c | 1.62 | 0.016400648 | Alpha-1-antitrypsin 1-3 | 2, 3 |
| P0CL69 | Zfp703 | 1.23 | 0.030611211 | Zinc finger protein 703 | 4, 5 |
| P33587 | Proc | 0.79 | 0.040787194 | Vitamin K-dependent protein C | 2, 4, 7 |
| Q03734 | Serpina3m | 0.75 | 0.019808496 | Serine protease inhibitor A3M | 3 |
| E9PVD2 | Itih4 | 0.58 | 0.007866917 | Inter alpha-trypsin inhibitor, heavy chain 4 | 1, 3 |
| A0A087WSN6 | Fn1 | 0.37 | 0.048648557 | Fibronectin | 1, 5, 7 |
| E9Q0S6 | Tns1 | -2.17 | 0.004727638 | Tensin 1 | 5 |
| G5E8B6 | Kirrel3 | -6.64 | 0.002307072 | Kin of IRRE-like protein 3 | 5, 6 |
